# Enriched pathogen diversity under host-type heterogeneity and immune-mediated competition

**DOI:** 10.1101/195065

**Authors:** Pamela P. Martinez, Robert J. Woods, Mercedes Pascual

## Abstract

Pathogen strains can stably coexist if they specialize on different hosts. Multiple strains can also coexist on a single host through negative frequency-dependent interactions mediated by partial cross-immunity. Understanding pathogen diversity remains a challenge however when both host specificity and cross-immunity are acting and may be functionally linked, as has been proposed for rotavirus, where a single protein is both antigenically important and determines host specificity by binding to the genetically encoded human blood group antigens. This situation is akin to the more general question in ecology of species coexistence when stabilizing and equalizing mechanisms interact. We examine this interaction with a theoretical model motivated by rotavirus and apply an adaptive dynamics framework to show how these two kinds of competition, typically considered separately, affect diversity. When cross-immunity depends on host-pathogen affinity, diversity is magnified as long-term evolution allows for the coexistence of multiple semi-specialized strains, similar to observations in rotavirus. In contrast, the simultaneous co-occurrence of several semi-specialized individuals is not observed when the degree of cross-immunity is independent from affinity distance among strains. The interplay of equalizing and stabilizing mechanisms fundamentally modifies diversity patterns and should be considered when addressing strain coexistence.

## Introduction

Motivated by the increased availability of molecular data, the interplay of population dynamics and molecular evolution is at the center of a better theoretical understanding of pathogen antigenic diversity (Grenfell et al. 2004, Lipsitch and O’Hagan 2007, Volz et al. 2013). Two kinds of phenotypic variation in pathogens influence this interplay, which concern respectively their differential ability to infect through binding to host receptors, and that to escape the immune system through antigenic diversity. Questions on strain diversity within pathogen populations are similar to those on species diversity within ecological communities. Specifically, two kinds of mechanisms for long-term species coexistence are distinguished in the ecological literature: those known as ‘stabilizing’ which operate in a frequency-dependent manner and confer an advantage to the rare; and those known as ‘equalizing’ because they minimize absolute fitness differences between species which would otherwise lead to dominance and exclusion (Chesson 2000). Equalizing effects reduce differences in fitness components such as growth rate, reproduction and survival. Given absolute fitness differences that promote exclusion, coexistence can also result from the existence of trade-offs as in the classical example provided by Tilman’s R* theory (Tilman 1982). Thus, multiple species that compete for the same resources can coexist if they have significantly different resource requirements and rely on the presence of a trade-off in the ability to consume these requirements.

In pathogens, a potential stabilizing mechanism is the frequency-dependent selection imposed by the immune system which can structure their populations into different coexisting strains (Gupta et al. 1996, Gupta et al. 1998, Gupta and Maiden 2001, Gog and Grenfell 2002, Gomes et al. 2002, Artzy-Randrup et al. 2012). The antigenic determinants recognized by the specific immune response are the phenotypic traits that underlie competition for hosts at the population level: once a host is protected against a given antigenic variant, it is no longer a resource for the growth of this specific type. Antigenic evolution allows the pathogen to escape immunity in the host population and strains that are sufficiently distant antigenically can co-exist by limiting similarity (Macarthur and Levins 1967). Coexistence arises here from purely stabilizing differences due to the frequency of given antigenic types, with no fixed fitness differences between individual pathogens. In contrast, equalizing mechanisms maintaining strain diversity in viruses concern fixed fitness differences and can arise from phenotypic variation in the host population itself. In particular, host variation has the potential to promote strain diversity if strains differ in their ability to bind host receptors. This is the case for example for both norovirus and rotavirus which recognize human histo-blood group antigens as receptors, with the ABO group and Lewis b antigen structures determining the strain’s specificity for the host (Shirato et al. 2008, Van Trang et al. 2014). Host types would therefore be akin here to resources in Tilman’s R* theory.

A more complete understanding of the structure of diversity within pathogen populations such as these requires therefore consideration of both stabilizing and equalizing forces operating together. We address here, with an adaptive dynamics approach, how differences in host-pathogen affinity (equalizing effect) interact with selection imposed by specific immunity (stabilizing effect) to influence strain diversity. Our results demonstrate the importance of a constraint imposed by a link between these two mechanisms for the outcome of this interaction and for the stable coexistence of pathogen strains. We discuss the relevance of these findings in the more general context of the ecology and evolution of infectious diseases.

## Methods

### One resident and one mutant

We start by implementing an extended *SIR* (Susceptive–Infected–Recovery) transmission model that considers two types of hosts, *A* and *B*, and a pathogen strain that differs in its affinity to bind each host receptor. This model is described by the following set of equations:

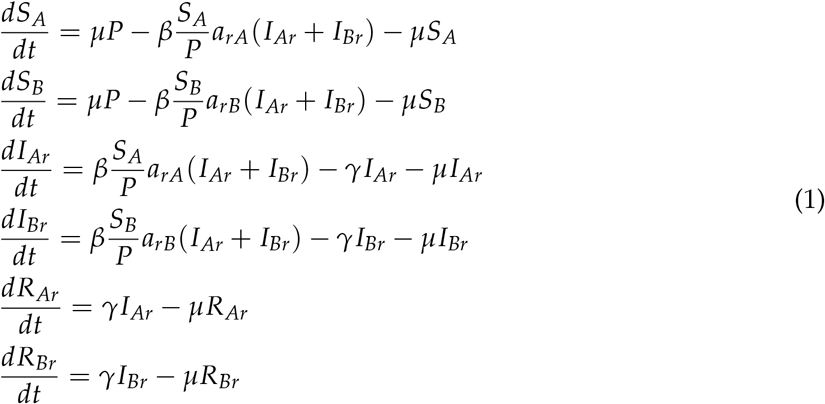

Within each host population, newly born individuals enter the susceptible class *S*. When hosts become infected, they move to the infected (and infectious) class *I*, and then to the recovery class *R* according to the recovery rate *γ*. The value of *γ* is considered constant because changes in this rate after repeated exposure have not been demonstrated in rotavirus. *P* refers to the population size of each host, and *μ*, to the birth/death rate (fig. 1A). For the purpose of this study, we have assumed that both host sub-populations are equally represented (*P* = *P*_*A*_ = *P*_*B*_).

**Figure 1:**
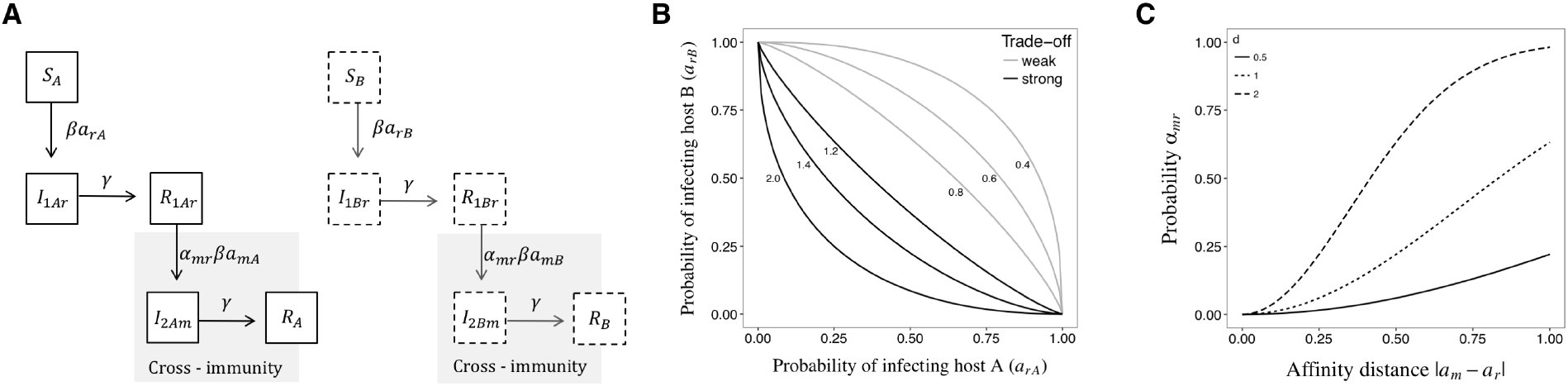
A) Transmission model. *S, I, R* refer respectively to the classes of Susceptible, Infected and Recovered individuals. The diagram on the left corresponds to host *A*, and the one on the right, to host *B*. B) Affinity trade-off. The trade-off can be strong (*s* > 1) or weak (*s* < 1). The different curves correspond to different values of s as indicated by the numbers next to each of them. C) Cross-immunity function. The probability *α* is plotted as a function of the affinity distance between current and previous infections, for different values of *d*. The epidemiological parameters that we have used are described in table A1.

We first consider the case where there is a single resident strain *r*. The force of infection consists of the contact rate *β* and the probability of being infected after contact *a_r_*, a value that varies between 0 and 1. Here, this probability *a_r_* specifically refers to host affinity and defines the identity of each strain. An increase in the affinity for one host is assumed to occur at the expense of a decreasing affinity for the other so that 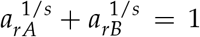 (Egas et al. 2004). This trade-off implies that each strain can be highly specialized for only one host, and in an extreme case, for example when *a_rA_* = 1, the strain *r* is completely specialized for of host *A*, and thus, *a*_*rB*_ = 0. The shape of this trade-off is determined by parameter *s*, where values of *s* > 1 result in a strong trade-off, while *s* < 1 generates a weak trade-off (fig. 1B). The endemic equilibrium for the system in equations (1) is given by:

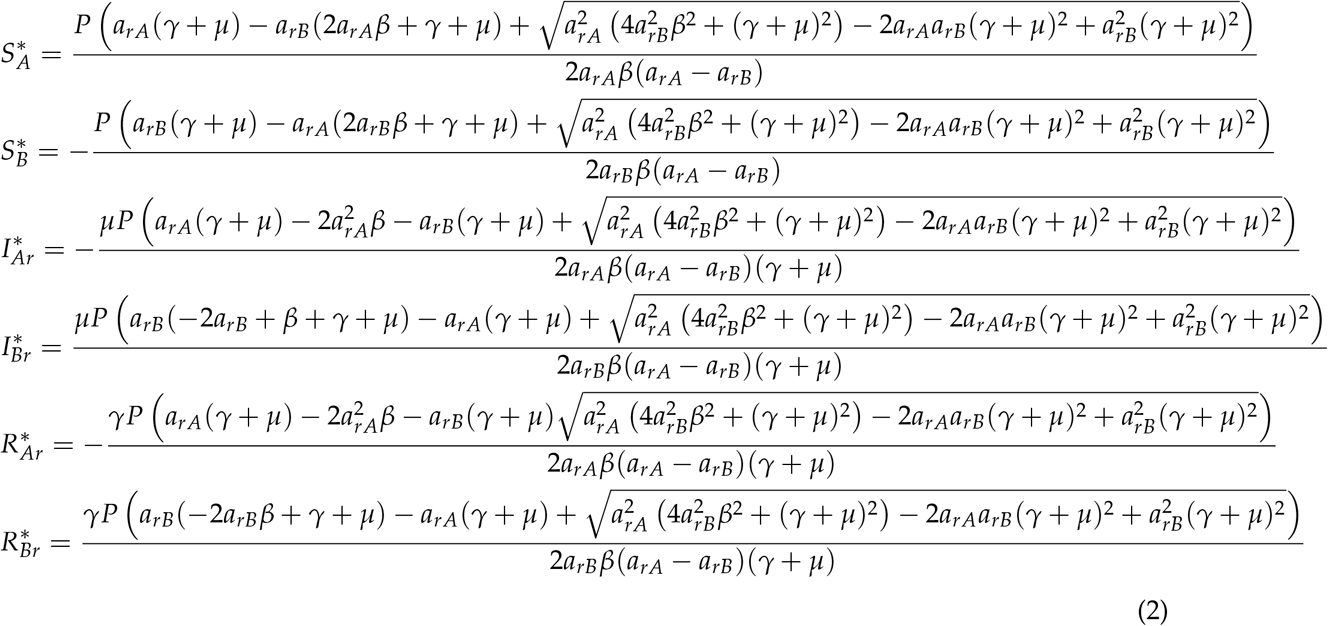

We then introduce a mutant strain *m* and ask under which conditions it will be able to invade the system at equilibrium. The new set of equations that characterizes population *A* is as follows:

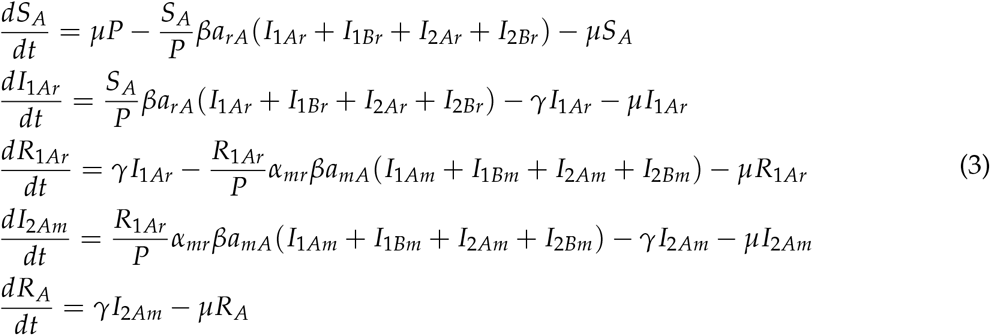

Similarly, the equations that characterize the population *B* are given by:

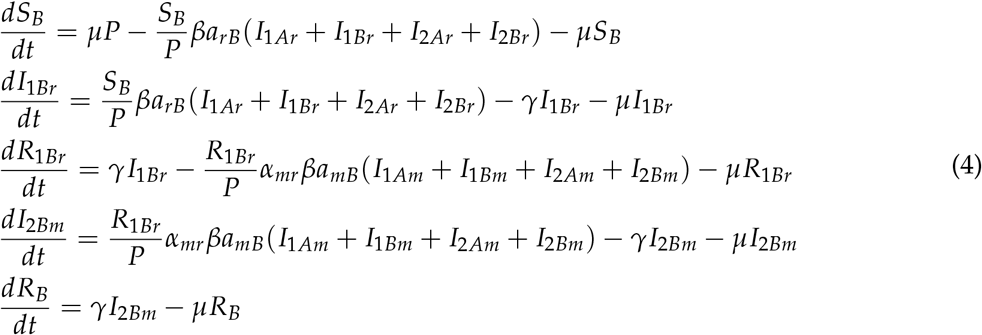

We analyze the evolution of the system via an adaptive dynamics framework (Metz et al. 1996, Dieckmann and Law 1996, Geritz et al. 1998), which overcomes some of the limitations of the classical approach based on maximizing *R*_0_, and allows in particular frequency-dependent pressures against common strains (Dieckmann 2002). This framework assumes that a mutant has a small difference in the phenotype of interest with respect to the resident strain. Changes in the growth rate associated with changes in the mutant phenotype determine the direction in which the population will evolve. Mutants can invade the system if the growth rate or invasion fitness *λ* is positive, which is defined by the leading eigenvalue of the Jacobian matrix of the system in the presence of the mutant (Metz et al. 1992). This matrix has a block-triangular form, where the diagonal blocks correspond respectively to the Jacobian of the resident *J_r_* (i.e. the system in the absence of the mutant) and the Jacobian of the mutant *J*_*m*_, as follows:

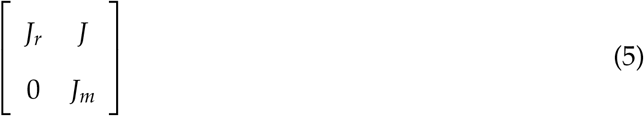

Thus, the eigenvalues of the new system are those of these two block-triangular matrices. Since a locally stable endemic equilibrium requires that the real part of the eigenvalue of *J*_*r*_ be negative, the stability of the system of equations (3,4) is determined by the leading eigenvalue of *J*_*m*_ which defines the invasion fitness *λ*. If the real part of *λ* is positive, the equilibrium becomes unstable and thus the population of the mutant will increase over time and invade the system; conversely, if *λ* is negative, the mutant will not be able to invade and will go extinct. The invasion fitness of a mutant *m* has the following form:

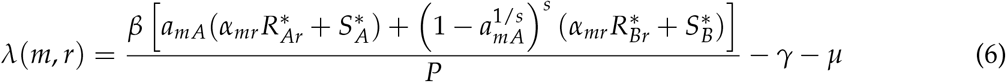

The direction of evolution for receptor binding specificity as the result of successive invasions is determined by the selection gradient *g*, which is given by the first derivative of *λ* with respect to the trait of the mutant, evaluated at the resident trait value:

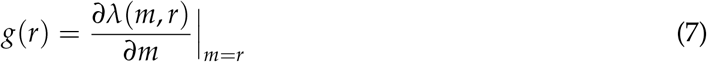

If the selection gradient is positive, mutants with affinity values greater than those of the residents will be successful, whereas if it is negative, mutants with lower trait values will invade. An evolutionary singular strategy *r\** is reached when *g*(*r*) = 0 (Geritz et al. 1998). The singular strategy is a convergence stable strategy when the first derivative of the selection gradient 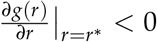, and it is an evolutionary repellor when it is greater than zero (Christiansen 1991). In addition, if the second derivative of the invasion fitness 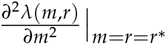 is negative, *r*\* is an evolutionary stable strategy (Smith 1982); whereas if it is positive, the singular strategy is an evolutionary branching point (Geritz et al. 1998). If the singular strategy is both convergent and evolutionary stable, then it is called Continuously Stable Strategy (CSS, Eshel 1983).

To compare the biological scenario where the degree of cross-immunity *α* is independent from the affinity to one in which these two traits are linked, we consider next two different models. The first model assumes that the values of *α* are independent from the identity of the strains, where *α* = 0 can be interpreted as complete cross-immunity, and *α* = 1 as a lack of cross-protection. The second model assumes that *α* depends on the distance between the affinity values of the current (*a*_*m*_) and past infection (*a*_*r*_), such that 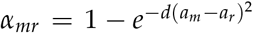, where the parameter *d* determines how fast immune protection decays as a function of the distance (fig. 1C, Gog and Grenfell 2002). This assumption implies that changes in the protein that binds the host receptor have an effect in both the degree of cross-immunity (antigenic distance), and the affinity for the host.

### Two or more residents, and one mutant

To expand the analysis to two or more residents, we rely on a numerical approach to analyze the possible evolutionary outcomes. For population *A* and *B*, the sets of equations are given by:

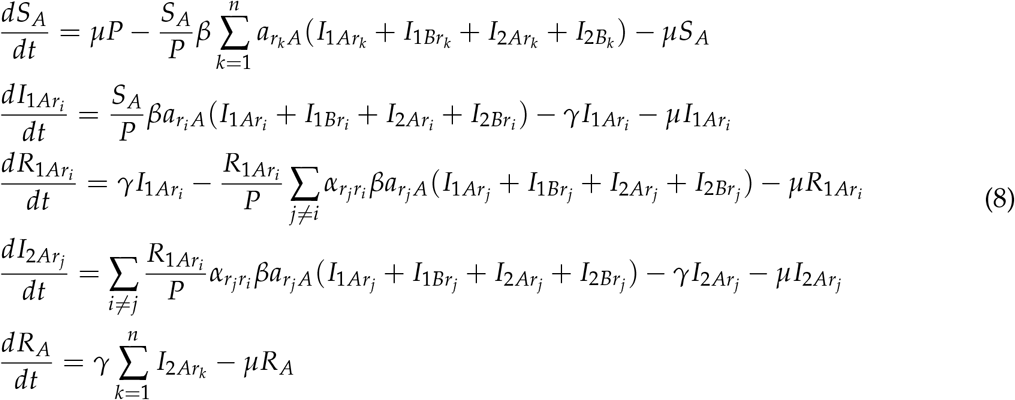

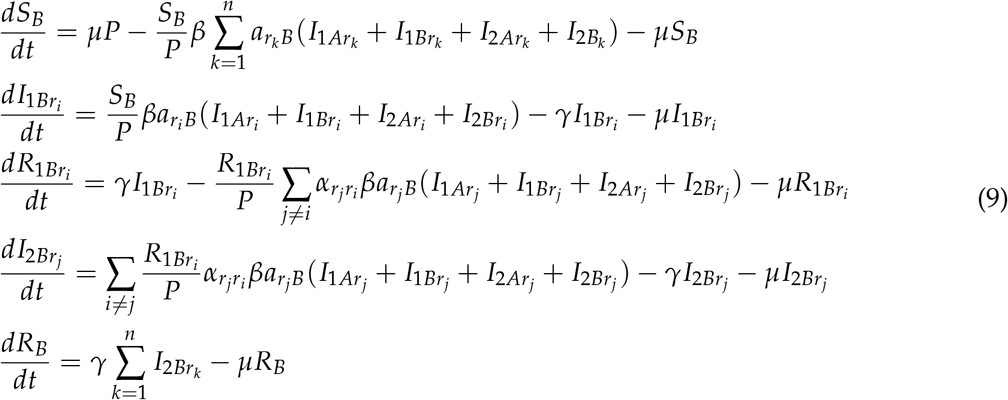

The first infection is represented by the strain *r*_*i*_ and its respective affinity for hosts *A* (*a_r_i_A_*) and *B* (*a_r_i_B_*). The second infection is then characterized by strain *r_j_*, where *r*_*i*_ and *r_j_* are two of *n* strains present at a given time. After the second infection, the host becomes immune to all future infections. The epidemiological parameters are described in table A1.

The invasion fitness of the mutant in the model with two or more residents is as follows:

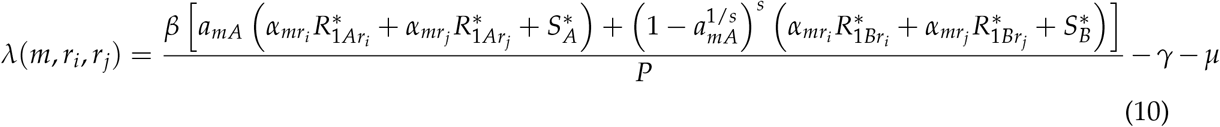

where *α*_*mr_i_*_ and *α*_*mr_j_*_ are either fixed parameters or a function of the respective distance between *a*_*m*_ and *a*_*r_i_*_, and between *a*_*m*_ and *a*_*r_j_*_. The direction of evolution as the result of successive invasions is determined by the selection gradient *g*, given by the first derivative of *λ* with respect to the trait of the mutant *m* and evaluated at the resident traits values *r*_*i*_ and *r*_*j*_:

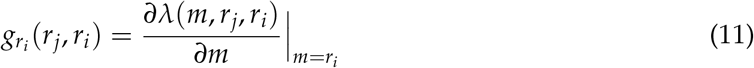

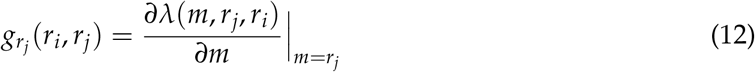

For this numerical implementation, we start all the simulations with a unique strain *r*, for which *a_rA_* = 0.5 and allow the system to evolve until it reaches a steady state (ecological time scale). We then introduce a mutant *m* from a normal distribution with mean equal to the value *a_rA_* in the ancestor strain and standard deviation of 0.001. We then repeat the inclusion of new mutants, one at a time, until the system reaches a steady state in the evolutionary time scale.

## Results

We first examine analytically a system consisting of one resident and one mutant strain under two conditions: an independent strength of cross-immunity *α*, and one in which *α* is linked to the host affinity distance. Figure 2A summaries the results for the model with independent *α*: under weak host affinity trade-offs (*s* < 1), the system evolves to a unique evolutionarily stable strategy (ESS), which is also a continuously stable strategy (CSS), over the full range of cross-immunity. Under this scenario, the strains evolve to have the same ability to infect both host types (*a*_*rA*_ = *a*_*rB*_), becoming neutral with respect to the binding affinity. When the trade-off is stronger (*s* > 1), evolutionary branching occurs, where the population evolves to a dimorphic population. Finally, under a very strong trade-off (*s* > 2), the most likely scenario is one in which both strains become specialized for the same host (evolutionary repellors), because even very strong cross-immunity cannot push one of the populations to utilize the other host due to the steep trade-off. Stable coexistence of two specialized strains could be possible, however, if the evolutionary steps were larger, as may be the case for zoonotic introductions, or large effect mutations. In summary, when *α* is fixed, a weak trade-off favors the evolution of generalist strains that are neutral with respect to their ability to bind the host receptor, while the evolutionary outcome for a strong trade-off is determined by initial conditions, where the two strains can become completely specialized for different hosts or for the same one. The results for the model in which the two phenotypes (differences in binding and immunity) are linked show substantial differences (fig. 2B). This assumption implies that changes in host-pathogen binding affinity have a direct effect on cross-protection given by the specific immunity acquired from past infections. Figure 2B is dominated by evolutionary branching points, followed by evolutionary repellors, and with only a small fraction of the parameter space ending in a continuously stable strategy, even for weak trade-offs (*s* < 1).

**Figure 2:**
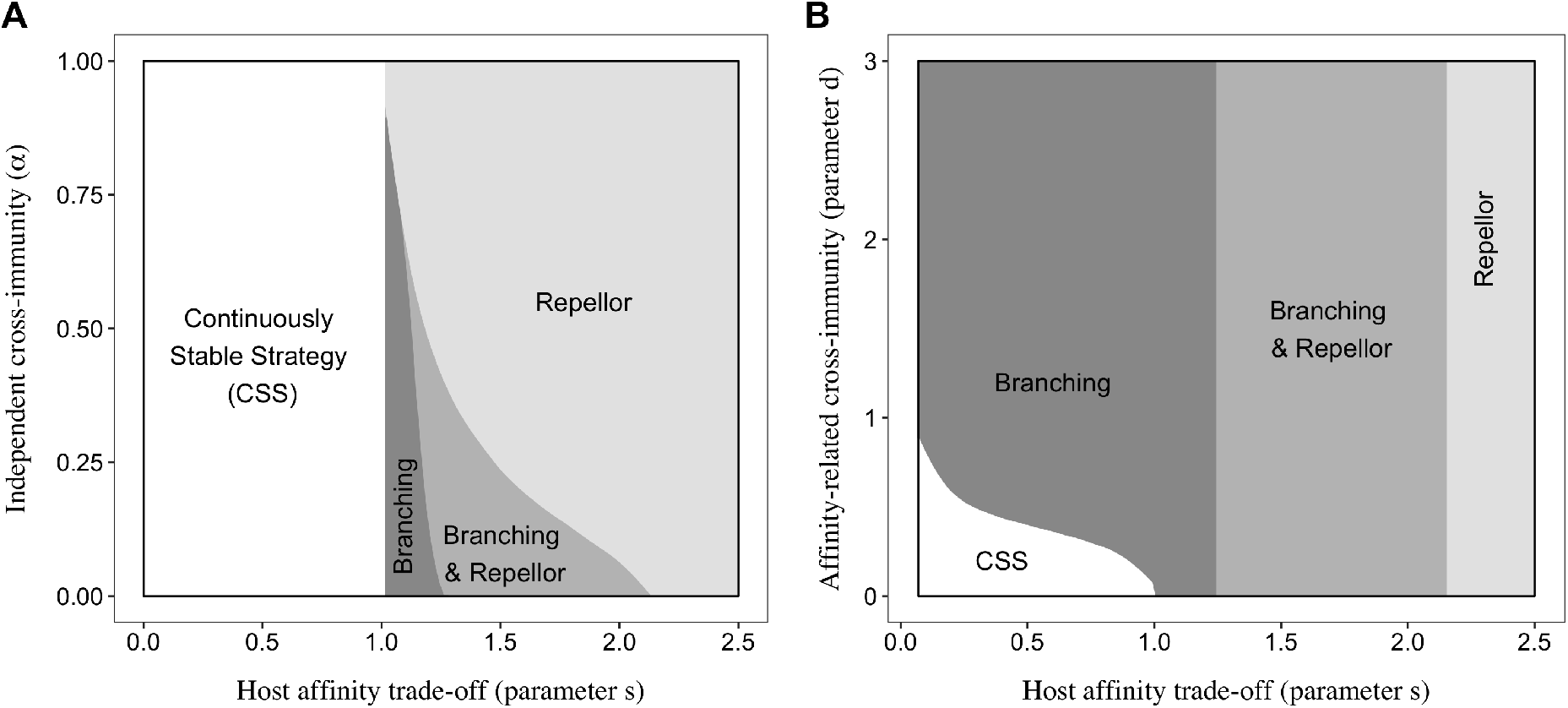
Evolutionary strategies as a function of the strength of cross-immunity and affinity trade-off. Parameter space under which the system evolves to a continuous stable strategy, evolutionary branching and/or repellor for independent (A) and affinity-related (B) values of cross-immunity, produced analytically. Complete cross-immunity is indicated by *α* = 0 and *d* = 0, while an absence of protection is reflected by *α* = 1 and very high values of *d*.

The analytical approach allowed us to establish the conditions under which two strains either stably coexist or exclude each other. However, to examine how the system behaves in the presence of two or more resident strains, we needed to implement a numerical approach. Concordant with our analytical results, we see that when the degree of cross-protection does not depend on the binding affinity, a weak trade-off leads to generalists, while a strong trade-off leads to full specialists, where each strain becomes completely specialized for the same or different hosts, avoiding competition (fig. 3A). By contrast, if the strength of cross-immunity is functionally related to the difference in host affinity, the dynamics of the system do not follow from the simpler models. In this case, after branching occurs, the population becomes polymorphic with completely or partially specialized pathogens, where the coexistence of multiple strains is now evolutionarily stable (fig. 3B, fig. A1). This result also indicates that higher values of *d*, and thus narrower cross-immunity width, amplify the coexistence of semi-specialized strains, which is consistent with observations in nature for pathogens like rotavirus, as we discuss later.

**Figure 3:**
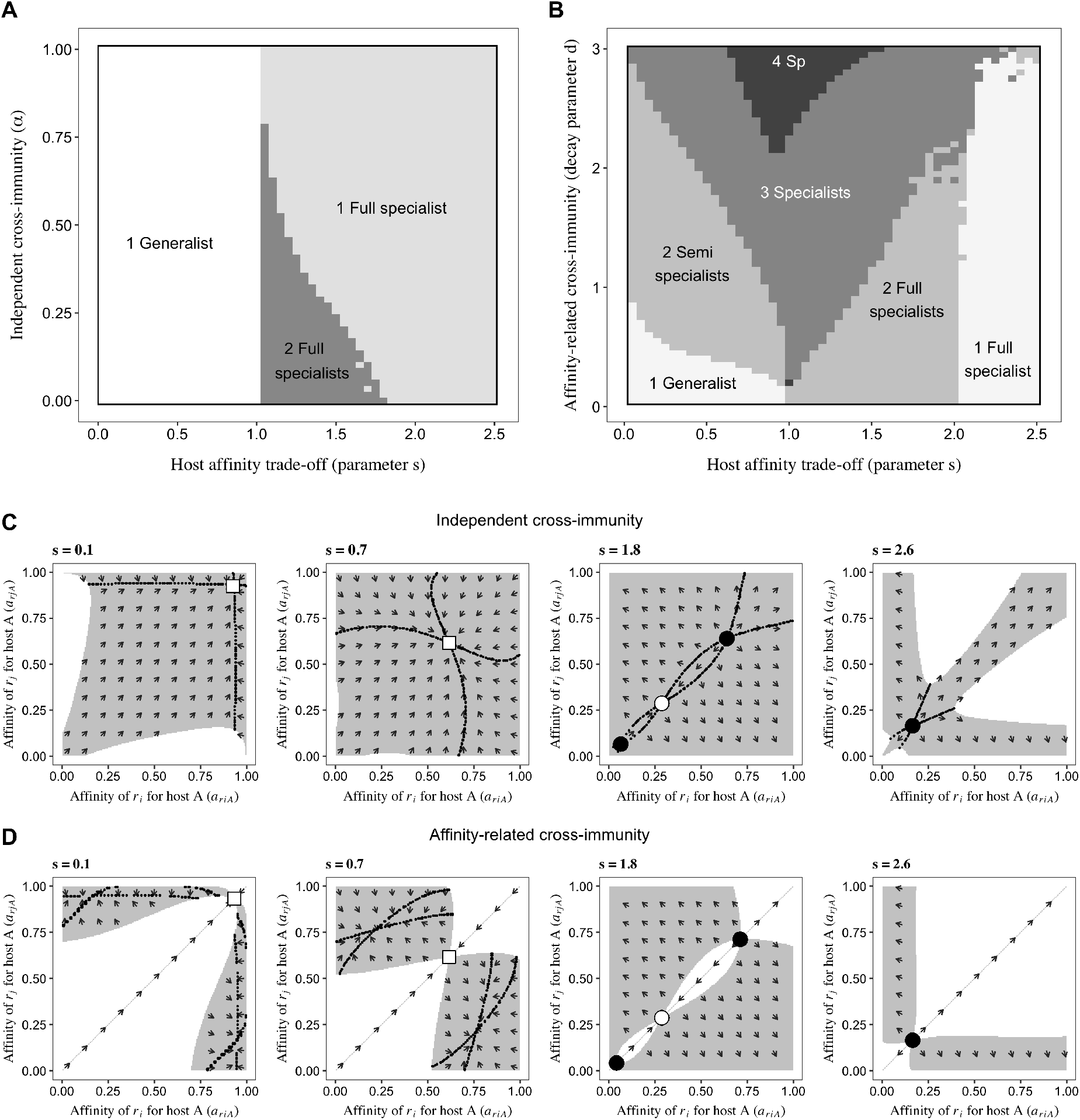
Long-term evolution. (A-B) Parameter space under which the system evolves to a generalist strain, full specialist, or to multiple semi specialists, for independent (A) and affinity-related (B) values of *α*. These results were numerically generated, with an initial affinity value *a*_*rA*_ = 0.5. (C-D) Trait evolution plots for the affinity for host A. Comparison of the evolutionary outcomes for the scenario in which *α_mr_* is fixed (*α* = 0.08, figure C) and the one in which it depends on the affinity distance between *r*_*i*_ and *r*_*j*_ (*d* = 0.5, figure D). The grey area indicates the region for which both traits *r_i_* and *r_j_* can mutually invade and coexist. The arrows indicate the direction of evolution as the result from successive invasion events, estimated from the selection gradients. Dotted black lines refer to the adaptive isoclines, where the selection gradient vanishes. The intersection of these isoclines denotes singular strategies, which are classified as CSS (white square), branching (white dot), or repellor (black dot).

Finally, we analyze the evolutionary dynamics of the system under the particular scenario of a high-degree of cross-immunity. Figure 3C shows the mutual invasibility plots under different assumptions about host type trade-offs for the model with independent cross-immunity. When there is a weak trade-off in host utilization (*s* < 1), we observe that the system evolves towards a singular generalist strategy. In contrast, for values of *s* > 1, the strains evolve to be specialists, either on different hosts (top left and bottom right of the mutual invasibility plot), or on the same host (bottom left and top right). The evolutionary outcome is dependent upon the initial values of the affinity trait, which can lead the system in the direction of an evolutionary branching point (attractors that then allow branching toward specialization on different hosts), and/or away from an unstable repellor, which can trap the strains in regions where they both specialize on the same host. As we vary the strength of the affinity trade-off, the evolutionary outcomes are summarized in the bifurcation diagram from figure A2A, which shows that the system is dominated by a CSS for values of *s* < 1. Likewise, for *s* ≥ 2 the only possible outcome is a repellor, and thus a completely specialized trait. From high to low values of *s*, a bifurcation occurs at *s* ≈ 2 where two repellors emerge together with a saddle node until s approaches 1. Figure 3D summarizes the findings for the affinity-related cross-immunity model under a broad strain space (*d* = 0.5), for which the average cross-immunity is analogous to *α* = 0.08. These results show that the coexistence of semi-specialized strains is possible, with the potential evolutionary outcomes shown in figure A2B. The area in the mutual invasibility plot under which two residents would coexist varies with the value of s in a nonlinear way (fig. A3); this area refers to the percentage of possible trait combinations under which the invasion fitness (growth rate) is positive for both strains, showing that intermediate values of the affinity trade-off allow for greater coexistence. Interestingly, a model with fixed *α* exhibits more transient ecological coexistence than the one with a variable cross-immunity, but this coexistence does not last when considering evolutionary time scales.

## Discussion

The model we have presented ties together the evolution of equalizing and stabilizing mechanisms. Previous studies of diversity in ecology have considered these two aspects of competitive dynamics independently. In the context of infectious diseases, coexistence of multiple pathogens has been addressed in the presence of different types of trade-offs, including the relationship between virulence and transmission rate (reviewed in Gandon 2004), and the emergence of specialization under habitat choice (i.e. host preference, Egas et al. 2004). Here we show that if the strength of cross-immunity (a stabilizing mechanism) is linked to the difference in host affinity (an equalizing mechanism), the evolutionary outcomes do not follow from the models when these are considered independent. Coexistence of multiple semi-specialized strains able to infect both types of hosts is the most common outcome, amplifying diversity. Our results also suggest that a weaker immune response is necessary to overcome the affinity trade-off (indicated by higher values of *d*, fig. 3B) to maintain 2 or more phenotypes coexisting. Concordant with previous theoretical studies, we also find that when the degree of specific immunity is independent from the affinity distance, a weak trade-off would lead to generalists, whereas a strong trade-off promotes the evolution of full specialists (Egas et al. 2004, Gudelj et al. 2004).

For pathogens such as rotavirus, both equalizing and stabilizing mechanisms are thought to be important. Several viral antigenic sites are located in the same subunits of the proteins responsible for binding the host-receptor (Taylor et al. 1987, Ruggeri and Greenberg 1991, Taylor and Dimmock 1994), suggesting that differences in each of these traits might be constrained to co-vary and could not occur independently as would be represented with simpler ecological models. The model that links equalizing and stabilizing mechanisms yields results consistent with the patterns of diversity seen in rotavirus. The number of variants in the protein that binds the host receptor has been shown to be higher than expected based solely on the number of histo-blood group antigens present in the human population (Matthijnssens et al. 2011). Additionally, strains do not appear to be completely specialized for particular host types (Van Trang et al. 2014, Ma et al. 2015). It would be valuable to further investigate whether viruses with significant diversity tend to use a range of similar receptors, or if less diverse viruses have become fixed on a single receptor with no similar receptors, as a consequence of different degrees of cross-immunity and trade-off. These studies would make possible the evaluation of predictions made by theoretical models like the one presented here.

Two of the main assumptions of the adaptive dynamics framework are that the system initially consists of a monomorphic population (i.e. all pathogens have the same trait), and that mutations have a small effect. These assumptions constrain the outcome when the direction of evolution is influenced by the presence of repellors that push the strains to specialize for the same host (trajectories along the diagonal in the trait evolution plots). However, many viruses can experience zoonotic introductions of somewhat different types, displacing the system into the region where two strains have very different affinity values. Alternative approaches to the analysis of the system have the potential to relax these limitations. For instance, the use of the Price equation would allow the relaxation of assumptions on small genetic changes. Recent studies have highlighted the generalization of this approach in the context of evolutionary dynamics (Nowak and Sigmund 2004, Day and Gandon 2006, Queller 2017, Lehtonen 2018). Future research is needed in this direction to assess whether results similar to ours would be obtained under this approach. In particular, mutation frequency and size should be further explored, together with the shape of the functional form connecting antigenic distance with cross-immunity, as this could affect the evolutionary dynamics in a nonlinear way.

Linking equalizing and stabilizing forces is an active area of research in the field of ecology. The recognition that these different kinds of traits underlie competition has helped structure theoretical and empirical studies of species diversity (Chesson 2000, Levine et al. 2009). However, extension of theory from two to multiple species remains limited (Barabás et al. 2018). The representation of equalizing and stabilizing forces as two-dimensional orthogonal axes has been useful to communicate their distinction (e.g. Adler et al. 2007, MacDougall et al. 2009, Mayfield and Levine 2010), but such graphical illustration implies independence that may not apply to the large diversity of phenotypic traits that motivates studies of coexistence in nature. Here we show that the manner in which these two processes are linked has an impact amount of the diversity and structure of that diversity. The non-independence of these axes and the underlying biological mechanisms they represent should be critically evaluated, as they have the potential to impact broad in ecology. When equalizing and stabilizing mechanisms are linked, we can expect very different outcomes for coexistence, with a potential amplification of overall diversity.

## Acknowledgments

We are grateful for the support of the Fogarty International Center at NIH [Program on the Ecology and Evolution of Infectious Diseases, EEID, R01-TW009670] to M.P., and the support provided by NIH K08 award 9(5K08AI119182-03) to R.J.W.

## Online Appendix

### A Supplementary Figures

**Table A1:** Epidemiological parameters*.

| Description (parameter) | Value |
| --- | --- |
| Birth/death rate ($\mu$) | 0.067 (1/year) |
| Recovery rate ($\gamma$) | 4 (1/year) |
| Contact rate ($\beta$) | 20 |

\* Parameter values were chosen based on the feasibility of steady states of the system for ecological time.

**Figure A1:**
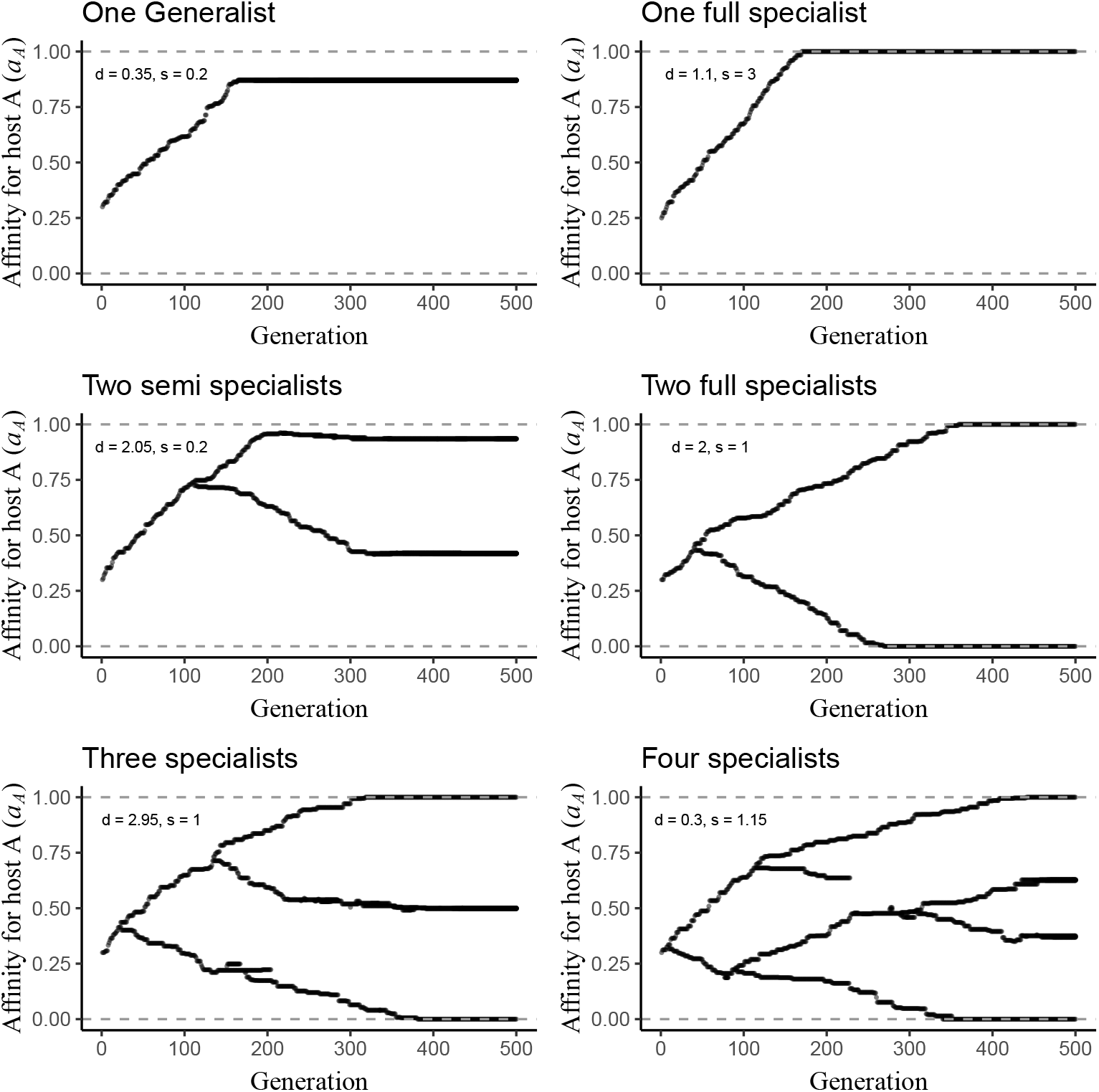
Representative evolutionary outcomes of affinity traits over time.

**Figure A2:**
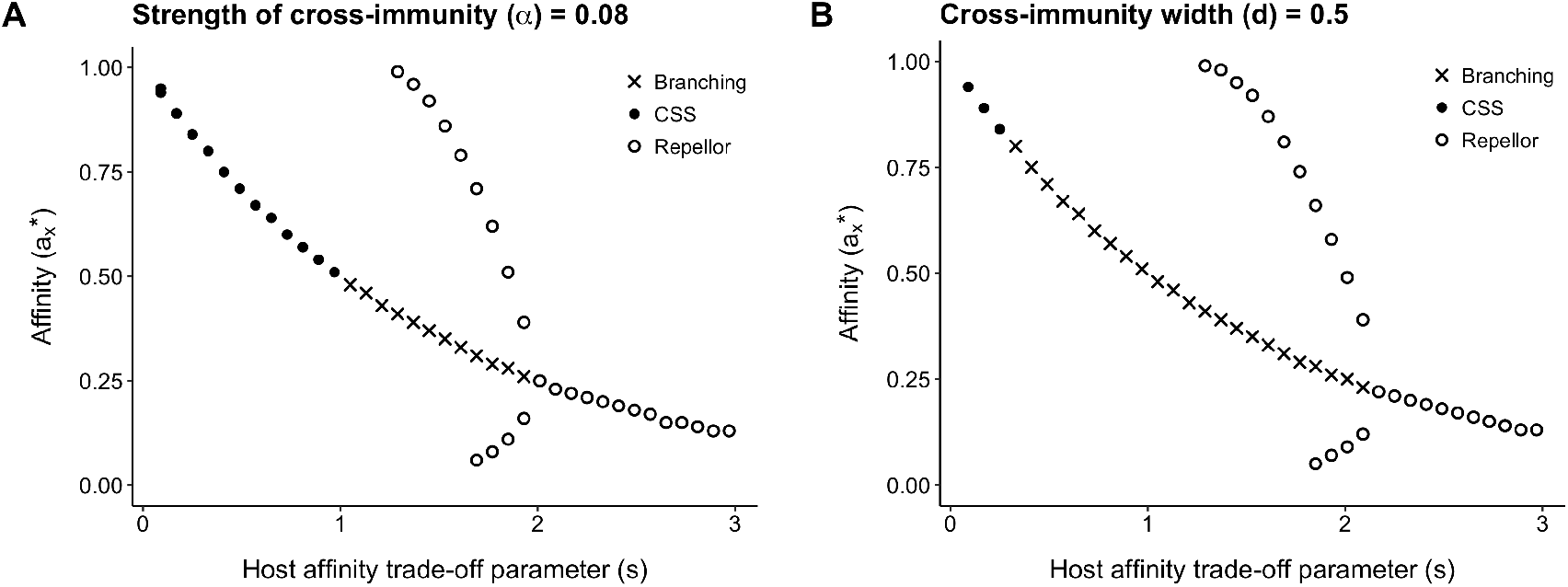
Bifurcation diagram. Evolutionary strategies as a function of the trade-off parameter *s*. The strategies under which the system may evolve are identified as: a Continuously Stable Strategy (CSS), a local evolutionarily stable strategy whose the affinity evolves until it reaches the CSS and cannot be invaded by a nearby mutant; an Evolutionary branching: the affinity trait evolves towards the branching point and once this point is reached, the traits diverge and the population becomes dimorphic; a Repellor: an evolutionary unstable strategy. The filled dots, open dots, and × symbols indicate CSS, repellors, and branching strategies, respectively, and the grey arrows denote the direction of monomorphic evolution.

**Figure A3:**
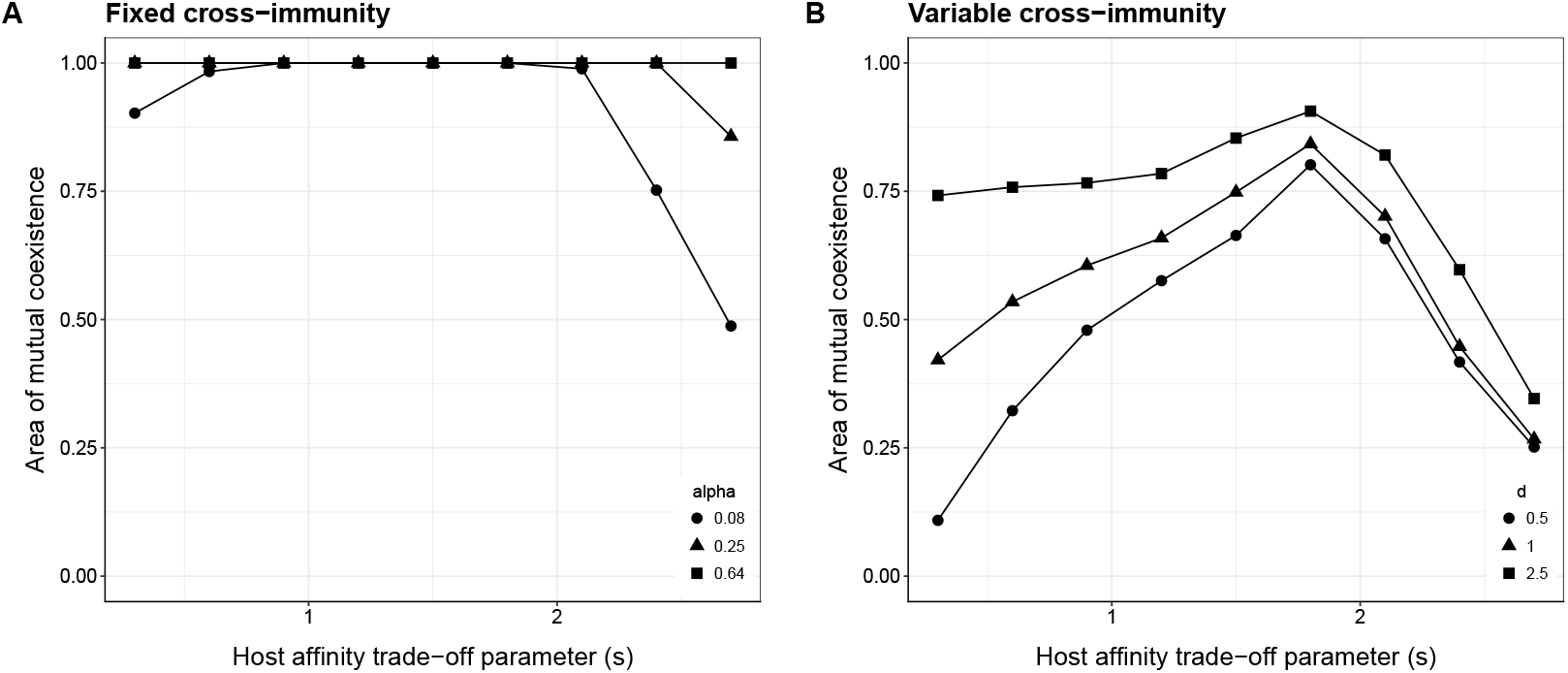
Percentage of all possible combinations of traits that allow for mutually invasion and coexistence as a function of *s*, for different values of fixed cross-immunity (A), and different values of *d* (variable cross-immunity, B).

## Literature Cited

Adler, P. B., J. HilleRisLambers, J. M. Levine. 2007. A niche for neutrality. Ecology letters 10(2): 95–104.

Artzy-Randrup, Y., M. M. Rorick, K. Day, C. Donald, A. P. Dobson, M. Pascual. 2012. Population structuring of multi-copy, antigen-encoding genes in Plasmodium falciparum. Elife 1: e00093.

Ayouni, S., K. Sdiri-Loulizi, A. de Rougemont, M. Estienney, K. Ambert-Balay, S. Aho, S. Hamami, M. Aouni, M. Neji-Guediche, P. Pothier, G. Belliot. 2015. Rotavirus P [8] infections in persons with secretor and nonsecretor phenotypes, Tunisia. Emerging infectious diseases 21(11): 2055–2058.

Barabás, G., R. D’Andrea, S. M. Stump. 2018. Chesson’s coexistence theory. Ecological Monographs 88(3): 277–303.

Chesson, P. 2000. Mechanisms of maintenance of species diversity. Annual review of Ecology and Systematics 31(1): 343–366.

Christiansen, F. B. 2000. On conditions for evolutionary stability for a continuously varying character. The American Naturalist 138(1): 37–50.

Day, T., G. Sylvain. 2006. Insights from Price’s equation into evolutionary. Disease evolution: models, concepts, and data analyses 71:23.

Dieckmann, U., R. Law. 1996. The dynamical theory of coevolution: a derivation from stochastic ecological processes. Journal of mathematical biology 34(5-6): 579–612

Dieckmann, U. 2002. Adaptive dynamics of pathogen-host interactions. IR-02-007

Eshel, I. 1983. Evolutionary and continuous stability. Journal of theoretical Biology 103(1): 99–111.

Egas, M., U. Dieckmann, M. W. Sabelis. 2004. Evolution restricts the coexistence of specialists and generalists: the role of trade-off structure. The American Naturalist 163(4): 518–531.

Gandon S. 2004. Evolution of multihost parasites. Evolution 58(3): 455–469.

Gentsch, J. R., A .R. Laird, B. Bielfelt, D. D. Griffin, K. Bányai, M. Ramachandran, V. Jain, N. A. Cunliffe, O. Nakagomi, C. D. Kirkwood, T. K. Fischer. 2005. Serotype diversity and reassortment between human and animal rotavirus strains: implications for rotavirus vaccine programs. Journal of Infectious Diseases 192(Supplement_1):S146–S159.

Geritz, S. A. H., E. Kisdi, G. Meszéna, J. A. J.. Metz. 1998. Evolutionarily singular strategies and the adaptive growth and branching of the evolutionary tree. Evolutionary ecology 12(1): 35–57.

Gog, J. R., B. T. Grenfell. 2002. Dynamics and selection of many-strain pathogens. Proceedings of the National Academy of Sciences 99(26): 17209–17214.

Gomes, M. G., G. F. Medley, D. J. Nokes. 2002. On the determinants of population structure in antigenically diverse pathogens Proceedings of the Royal Society of London B: Biological Sciences 269(1488): 227–233.

Grenfell, B. T., O. G. Pybus, J. R. Gog, J. L. N. Wood, J. M. Daly, J. A. Mumford, et al. 2004. Unifying the epidemiological and evolutionary dynamics of pathogens. Science 303(5656): 327–332.

Gudelj I., F. Van den Bosch, C. A. Gilligan. 2004. Transmission rates and adaptive evolution of pathogens in sympatric heterogeneous plant populations. Proceedings of the Royal Society of London B: Biological Sciences 271(1553): 2187–2194.

Gupta, S., M. C. J, Maiden, I. M. Feavers, S. Nee, R. M. May, R. M. Anderson. 1996. The maintenance of strain structure in populations of recombining infectious agents. Nature medicine 2(4): 437–442.

Gupta, S., N. Ferguson, R. Anderson. 1998. Chaos, persistence, and evolution of strain structure in antigenically diverse infectious agents. Science 280(5365): 912–915.

Gupta, S., M. C. J, Maiden. 2001 Exploring the evolution of diversity in pathogen populations. Trends in microbiology 9(4): 181–185

Kirkwood, C. D., R. F. Bishop, B. S. Coulson. 1996. Human rotavirus VP4 contains strain-specific, serotype-specific and cross-reactive neutralization sites. Archives of virology 141(3): 587–600

Larralde, G. I., B. G. Li, A. Z. Kapikian, M. A. Gorziglia. 1991. Serotype-specific epitope (s) present on the VP8 subunit of rotavirus VP4 protein. Journal of virology 65(6): 3213–3218

Lehtonen, J. 2018. The price equation, gradient dynamics, and continuous trait game theory. The American Naturalist 191(1): 146–153.

Levine, J. M., J. HilleRisLambers. 2009. The importance of niches for the maintenance of species diversity. Nature 461(7261): 254.

Lipsitch M., J. J O’Hagan. 2007. Patterns of antigenic diversity and the mechanisms that maintain them. Journal of the Royal Society Interface 4(16): 787–802.

Liu Y., T. A. Ramelot, P. Huang, Y. Liu, Z. Li, T. Feizi., W. Zhong, F. T. Wu, M. Tan, M. A. Kennedy, X. Jiang. 2016. Glycan Specificity of P [19] Rotavirus and Comparison with Those of Related P Genotypes. Journal of virology 90(21): 9983–9996.

Ma X., X. Sun, Y. Guo, J. Xiang, W. Wang, L. Zhang, Q. Gu, Z. Duan. 2015. Binding patterns of rotavirus genotypes P [4], P [6], and P [8] in China with histo-blood group antigens. PloS one 10(8): e0134584.

Macarthur, R., R. Levins. 1967. The limiting similarity, convergence, and divergence of coexisting species. The American Naturalist 101(921): 377–385

MacDougall, A. S., B. Gilbert, J. M. Levine. 2009. Plant invasions and the niche. Journal of Ecology 97(4): 609–615.

Matthijnssens J., M. Ciarlet, S. M. McDonald, H. Attoui, K. Bányai, J. R. Brister, J. Buesa, M. D. Esona, M. K. Estes, J. R. Gentsch, M. Iturriza–Gómara M. 2011. Uniformity of rotavirus strain nomenclature proposed by the Rotavirus Classification Working Group (RCWG). Archives of virology 156(8):1397–1413.

Mayfield M. M, J. M. Levine. 2010. Opposing effects of competitive exclusion on the phylogenetic structure of communities. Ecology letters 13(9): 1085–1093.

Metz, J. A., R. M. Nisbet, S. A. Geritz. 1992 Adaptive dynamics, a geometrical study of the consequences of nearly faithful reproduction Trends in Ecology & Evolution 7(6):198–202

Metz, J. A., S. A. Geritz, G. Meszéna, F. J. Jacobs, J. S. Van Heerwaarden. 1996 Adaptive dynamics, a geometrical study of the consequences of nearly faithful reproduction Stochastic and spatial structures of dynamical systems 45:183–231

Ndifon W., N. S. Wingreen, S. A. Levin. 2009. Differential neutralization efficiency of hemagglutinin epitopes, antibody interference, and the design of influenza vaccines. Proceedings of the National Academy of Sciences 106(21): 8701–8706.

Nowak, M. A., K. Sigmund. 2004. Evolutionary dynamics of biological games. Science 303(5659): 793–799.

Nordgren J., S. Sharma, F. Bucardo, W. Nasir, G. Günaydm, D. Ouermi, L. W. Nitiema, S. Becker-Dreps, J. Simpore, L. Hammarström, G. Larson. 2014. Both Lewis and secretor status mediate susceptibility to rotavirus infections in a rotavirus genotype–dependent manner. Clinical Infectious Diseases 59(11):1567–1573.

Queller D. C. 2017. Fundamental theorems of evolution. The American Naturalist 189(4): 345–353.

Ramani S., L. Hu, B. V. V. Prasad, M. K. Estes. 2016. Diversity in Rotavirus–Host Glycan Interactions: A “Sweet” Spectrum. CMGH Cellular and Molecular Gastroenterology and Hepatology 2(3): 263–273.

Ruggeri F. M., H. B. Greenberg. 1991 Antibodies to the trypsin cleavage peptide VP8 neutralize rotavirus by inhibiting binding of virions to target cells in culture. Journal of virology 65(5): 2211–2219.

Shirato, H., S. Ogawa, H. Ito, T. Sato, A. Kameyama, H. Narimatsu, Z. Xiaofan, T. Miyamura, T. Wakita, K. Ishii, N. Takeda. 2008. Noroviruses distinguish between type 1 and type 2 histo-blood group antigens for binding. Journal of virology 82(21): 10756–10767.

Smith, J. M. 1982. Evolution and the Theory of Games. Cambridge Univ. Press.

Sun X., N. Guo, D. Li, M. Jin, Y. Zhou, G. Xie, L. Pang, Q. Zhang, Y. Cao, Z. Duan. 2016a. Binding specificity of P [8] VP8* proteins of rotavirus vaccine strains with histo-blood group antigens. Virology 495:129–135.

Sun X., D. Li, R. Peng, N. Guo, M. Jin, Y. Zhou, G. Xie, L. Pang, Q. Zhang, J. Qi, Z. Duan. 2016b. Functional and structural characterization of P [19] rotavirus VP8* interaction with histo-blood group antigens. Journal of Virology 90(21):9758–9765.

Sun X., N. Guo, J. Li, X. Yan, Z. He, D. Li, M. Jin, G. Xie, L. Pang, Q. Zhang, N. Liu. 2016c. Rotavirus infection and histo-blood group antigens in the children hospitalized with diarrhoea in China. Clinical Microbiology and Infection 22(8):740–e1.

Taylor H. P., S. J. Armstrong, N. J. Dimmock. 1987. Quantitative relationships between an influenza virus and neutralizing antibody. Virology 159:288–298.

Taylor H. P., N. J. Dimmock. 1994 Competitive binding of neutralizing monoclonal and polyclonal IgG to the HA of influenza A virions in solution: Only one IgG molecule is bound per HA trimer regardless of the specificity of the competitor. Virology 205:360–363.

Tilman, D. 1982. Resource competition and community structure. Princeton university press.

Van Trang, N., H. T. Vu, N. T. Lee, P. Huang, X. Jiang, D. D. Anh. 2014. Association between norovirus and rotavirus infection and histo-blood group antigen types in Vietnamese children. Journal of clinical microbiology 52(5): 1366–1374.

Volz, E. M., K. Koelle, T. Bedford. 2013. Viral phylodynamics. PLoS Comput Biol 9(3): e1002947.

Zhang X., Y. Long, M. Tan, T. Zhang, Q. Huang, X. Jiang, W. Tan, J. Li, G. Hu, S. Tang, Y. Dai. 2016. P [8] and P [4] Rotavirus Infection Associated with Secretor Phenotypes Among Children in South China. Scientific Reports 6:34591.

